# Temporal and Spectral Neural Complexity Reveal Graded Auditory Awareness

**DOI:** 10.64898/2026.04.20.719685

**Authors:** Alberto Liardi, Daniel Bor, Fernando E. Rosas, Pedro A.M. Mediano

## Abstract

Recent advances have shown that the complexity of neural signals tracks global states of consciousness, such as wakefulness versus sleep. However, it is still unclear to what extent neural complexity reflects fine-grained changes in conscious content within the same global state. Here, we investigate how the complexity of brain signals is affected by increased perceptual clarity of a stimulus. To this end, we estimated neural signal complexity using Complexity via State-space Entropy Rate (CSER) to EEG recordings from an auditory discrimination task. In this paradigm, auditory stimuli were presented at varying signal-to-noise ratios (SNRs), with higher SNRs corresponding to greater subjective audibility and perceptual clarity, enabling us to relate neural complexity to graded perceptual awareness within a constant global state of consciousness. Our results showed that, while broadband CSER remains constant across SNRs, its spectral decomposition displays frequency-specific effects, with higher SNRs associated with a decreased complexity in *α* and *β* bands, increased complexity in *δ*, and no significant changes in *γ*. Additionally, a temporal investigation of CSER exhibited a significant increase in complexity with stimulus clarity, with deviations from baseline peaking approximately 30 ms before the ERP. Extending this analysis to pairs of brain regions, mutual information rate uncovered a sudden post-stimulus breakdown in long-range information transmission relative to baseline. Taken together, these results reveal that while aggregated complexity measures track global states of consciousness, time- and frequency-resolved information-theoretic measures can capture variations in perceptual awareness, demonstrating their sensitivity as estimators of the level of conscious experience.

## I. INTRODUCTION

In recent years, much progress has been made in determining the neural correlates of consciousness (NCCs) – the minimal set of neural mechanisms jointly sufficient for any given conscious experience [1–8]. Among the many approaches to finding NCCs, one of the most successful has been the analysis of neural complexity [9– 12], making complexity measures a central tool for probing different levels of consciousness across a wide range of conditions [13]. Studies have demonstrated the ability of these methods to distinguish disorders of consciousness [14], depth of anaesthesia [15, 16], and sleep stages [17, 18]. Increased complexity has also been observed under psychedelics [19–22], while consistent changes in complexity have been linked to psychiatric and neurological conditions such as depression [23–25] and schizophrenia [22, 23, 26, 27]. Beyond neuropathology, complexity has further been studied in meditation [28, 29], in the ageing of infants and adults [30–33], and in many other contexts and conditions [34–42].

While the above body of work has focused on global *states* of consciousness, another line of research has investigated whether neural complexity can also track the *contents* of consciousness and temporal fluctuations in awareness. Although less explored, existing studies provide preliminary evidence: complexity increases during emotional processing [43], correlates with mental effort in cognitive tasks [12, 44–46], and differentiates states of mind-wandering [47–51]. However, these findings are often based on stationary estimates of complexity, limiting the sensitivity of these measures to the fast, stimulus-locked dynamics of conscious perception. This is particularly relevant for task-related paradigms, where neural activity is inherently stimulus-evoked and non-stationary, and standard complexity measures might lack the temporal and spectral resolution required to characterise such dynamics. As a result, it remains unclear how neural complexity relates to changes in perceptual awareness and whether it can reflect transient, frequency-specific modulations tied to stimulus processing.

To address this gap, the present study analyses EEG data from an auditory discrimination task [52] to explore how neural complexity and information transmission are modulated over time and space by the perceptual awareness of the acoustic stimulus. In this experiment, participants are presented with auditory cues embedded in noise at varying signal-to-noise ratios (SNRs), ranging from indistinguishable (low SNRs) to clearly perceptible (high SNRs). Hence, this design provides a controlled framework to examine the relationship between information-theoretic measures of neural activity and graded levels of perceptual awareness within a constant global state of consciousness. We conduct our analysis using *Complexity via State-space Entropy Rate* (CSER) [21], an entropy rate-based complexity measure that, unlike traditional estimators employed in previous studies, provides a temporally and spectrally resolved quantification of the signal complexity. By comparing the neural activity across SNR levels – that is, by systematically varying degrees of stimulus clarity – we then investigate how perceptual awareness modulates neural complexity across time, space, and frequency bands. In parallel, we use the mutual information rate (MIR) – which quantifies the information transmitted between pairs of brain regions, or equivalently, the entropy rate they share – to extend this framework to large-scale inter-regional dynamics, inspecting how stimulus-dependent changes relate to the spatio-temporal structure of information transmission across the brain. Together, these measures enable a fine-grained investigation of how perceptual awareness shapes the structure and flow of entropy and information in ongoing brain activity.

Our results show that, contrary to studies of global states of consciousness, the temporal and spectral dimensions of CSER are necessary to discriminate perceptual levels, as global broadband complexity remains unchanged across SNRs. Concurrently, MIR exhibits a post-stimulus breakdown in shared entropy rate across cortical regions, suggesting a rapid reorganisation of information processing when the auditory stimulus is most clearly perceived. With these insights, the present work moves beyond the traditional use of complexity as a coarse marker of global conscious state, instead characterising its role in perceptual processing dynamics. In doing so, it contributes to the development of objective, information-theoretic markers of awareness that can be robustly estimated from non-invasive neuroimaging data in both healthy and clinical populations.

## II. RESULTS

We analysed EEG data originally reported by Sergent *et al*. [52]. The experiment involved auditory stimuli (French vowels “a” and “e”) presented at five signal-to-noise ratios (SNR) (–13 to *−*5 dB) plus noise-only trials, with participants being asked to identify the vowels and rate their audibility. Lower SNRs rendered the stimuli difficult or impossible to consciously perceive, resulting in reduced identification accuracy and lower audibility ratings, whereas higher SNRs increased perceptual clarity and reported awareness. In the original study, the authors identified a threshold for conscious access at SNR = *−*9 dB, marking the transition between predominantly unconscious and consciously perceived trials [52]. Data from 20 adult participants were recorded using 64 channels at a sampling frequency of 500 Hz, and each condition included 160 trials, totalling 960 trials per session. Further information about the dataset is reported in Sec. IV D.

To quantify the complexity of neural dynamics, we use Complexity via State-space Entropy Rate [21]. CSER measures the unpredictability of a time series by estimating the entropy rate of a state-space (SS) model fit to the signal. This captures the entropy of the process’s components that cannot be predicted from its past, providing a principled way to quantify how much new information is introduced as the system’s evolution unfolds over time [21]. We consider two complementary versions of the measure. *Steady-state* CSER provides a scalar estimate of signal complexity, reflecting the overall unpredictability of the process and being tightly linked to the commonly used Lempel-Ziv complexity [21, 53]. A powerful feature of steady-state CSER is that it can be decomposed into specific frequencies, yielding insights into how complexity is modulated by slow and fast oscillations. Alternatively, *time-resolved* CSER tracks how complexity evolves over time, offering a dynamic view of changes in neural activity [21]. Together, these measures provide insights into both the global and local structure of neural complexity. A complete description of the methods can be found in Sec. IV B, with schematic representations of the estimators shown in Figs. 1a-1b and 2a.

**FIG. 1:**
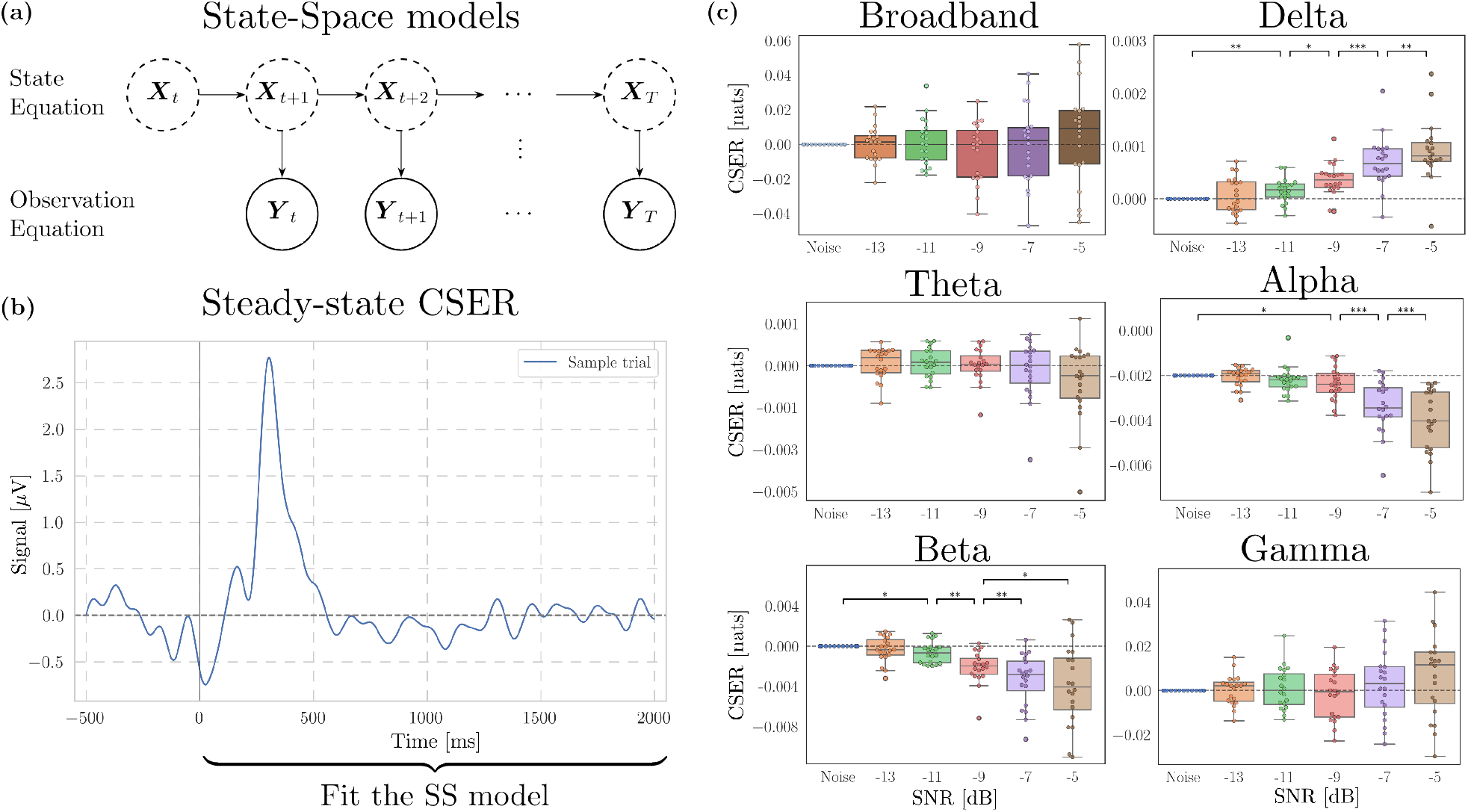
Band-specific complexity shows significant changes in perceptual clarity levels while broadband complexity remains unaffected. **(a)** CSER is based on state-space models, which estimate latent dynamics driving observable data. **(b)** We calculate steady-state CSER by fitting a single SS model to the full post-stimulus period. **(c)** Broadband and frequency-decomposed steady-state CSER for different SNR levels and for all subjects. Low SNR levels indicate weak or absent conscious perception, whereas higher SNRs suggest clearer and more reliably perceived stimuli. The CSER values shown are the differences w.r.t. CSER of pure noise. Only pairwise t-tests between adjacent SNRs are shown – the full statistical analysis is reported in the Supplementary Material. (P-values calculated with a one-sample t-test against the zero-mean null hypothesis. *: *p <* 0.05; **: *p <* 0.01; ***: *p <* 0.001).

**FIG. 2:**
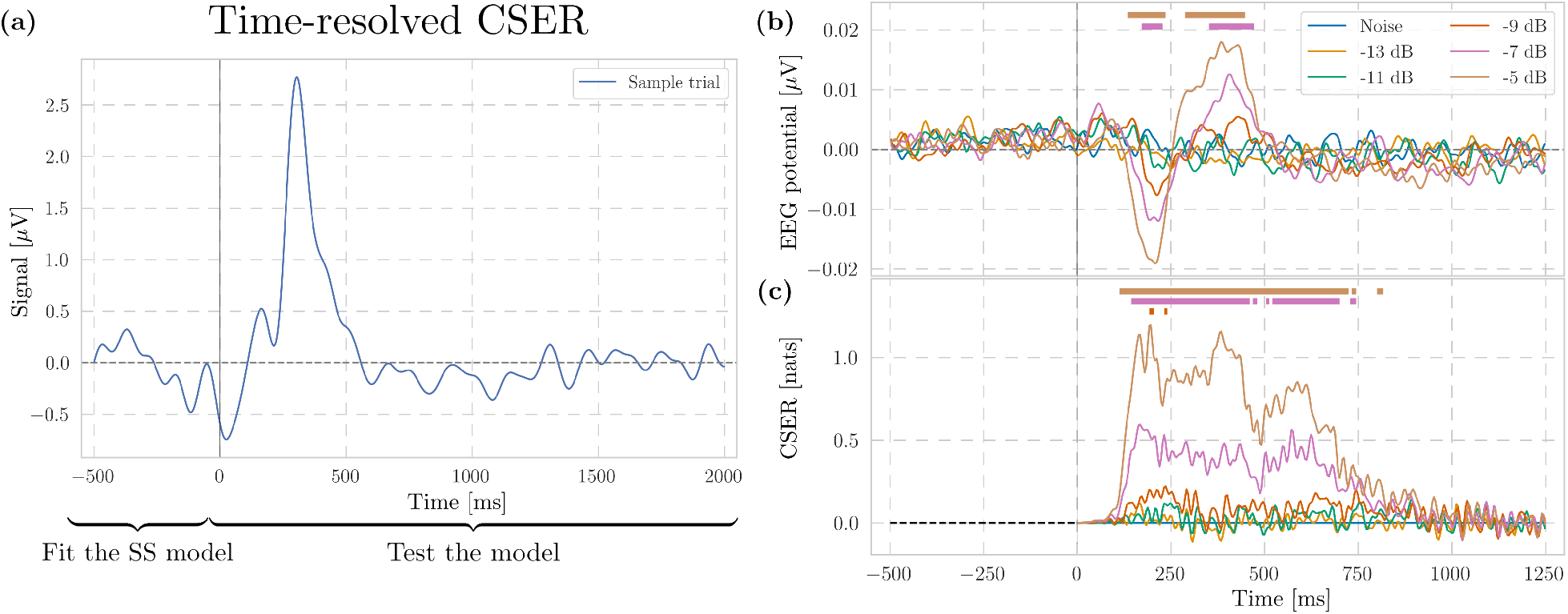
Time-resolved CSER discriminates between perceptual levels, with increased complexity for higher SNRs. **(a)** We calculate time-resolved CSER by fitting a state-space model on pre-stimulus data and evaluating it post-stimulus. **(b)** EEG ERP grouped by SNR levels averaged across all channels and subjects. **(c)** Time-resolved CSER grouped by SNR levels and averaged across all channels and subjects. CSER values shown are the differences w.r.t. CSER of pure noise. Coloured horizontal bars indicate significance level with *p <* 0.05 with a paired cluster permutation test against noise. For visualisation purposes, CSER and EEG signals above were low-passed to 50 Hz. The full statistical analysis is reported in the Supplementary Material.

### A. Steady-state and spectrally resolved neural complexity

We begin our analyses by focusing on steady-state CSER, which corresponds to the “average” complexity of the neural signal over time. We computed it by first fitting a state-space model on the signal post-ERP across all channels, and then estimated it for each subject and signal-to-noise ratio level (Fig. 1b). Interestingly, the broadband measure is not able to differentiate the various SNRs (Fig. 1c), not even between the clearest sound (SNR *−*5 dB) and pure noise (Noise). An additional spatial analysis did not show any regional difference (Fig. 6 Supplementary Material).

However, if we decompose CSER into frequency bands, we observe that higher SNRs are characterised by larger CSER in *δ* (*p <* 0.01 under repeated-measures ANOVA and Holm-Bonferroni corrected pairwise t-test), and lower in *α* (*p <* 0.001) and *β* (*p <* 0.05) (Fig. 1c). Interestingly, *α* band only discriminates between consciously perceived stimuli (SNR −9, −7, *−*5 dB) and noisier ones (SNR Noise, −13, *−*11 dB). In contrast, the *β* and *δ* bands also distinguish SNR *−*11 dB – i.e., just below the conscious threshold [52] – from Noise and SNR *−*13 dB. Finally, the complexity of the *γ* band (30 *−*100 Hz) is one order of magnitude larger than the others, thus being the main driver of the lack of significant differences in the broadband signal.

### B. Time-resolved neural complexity

For the study of time-resolved CSER, we first inspected the shape of the average event-related potential (ERP) for different SNR levels (Fig. 2b). This shows marked differences (*p <* 0.01 under ANOVA cluster permutation test [54] and Holm-Bonferroni corrected post-hoc pairwise t-test) between clearly consciously perceived stimuli (SNR *−*7, *−*5 dB) and noisier sounds (Noise, *−*9, *−*11, *−*13 dB) in the time ranges from 175 to 225 ms and from 375 to 450 ms. We then applied time-resolved CSER (see Sec. IV E) to obtain the evolution of complexity over time (Fig. 2c). Time-resolved CSER shows pronounced peaks around 200, 400, and 500 ms, corresponding to the first dip and the two peaks in the ERP. Repeated-measures ANOVA and pairwise cluster permutation tests show significant differences between SNRs *−*7 and *−*5 dB versus all the other SNRs at various timepoints, broadly between 150 and 750 ms (Supplementary Material, Fig. 8). Interestingly, the initial CSER peak precedes the corresponding ERP component by approximately 30 ms, replicating an analogous finding obtained in the original CSER study [21].

Next, we compared how time-resolved CSER is spatially distributed across brain areas (Fig. 3). Focusing on SNR *−*5 dB (the condition with highest stimulus clarity), we divided the 64 channels into 5 brain regions – frontal, temporal, central, parietal, and occipital (see Tab.I in Supplementary Material) – and for each of these we computed the spatial average of time-resolved CSER across all subjects and channels in that region (Fig. 3a). We observe that the temporal evolution of complexity differs across brain regions. A cluster-based permutation ANOVA (*p <* 0.01) followed by Bonferroni-corrected pairwise t-tests (*p <* 0.001) revealed a significant occipital cluster peaking around 300 ms, with other regions showing no statistically significant differences. Moreover, the single-channel activity shows substantial variability even within each brain area (Fig. 3b). In particular, the central region exhibits peaked activity at 300 ms, though this does not survive statistical testing.

**FIG. 3:**
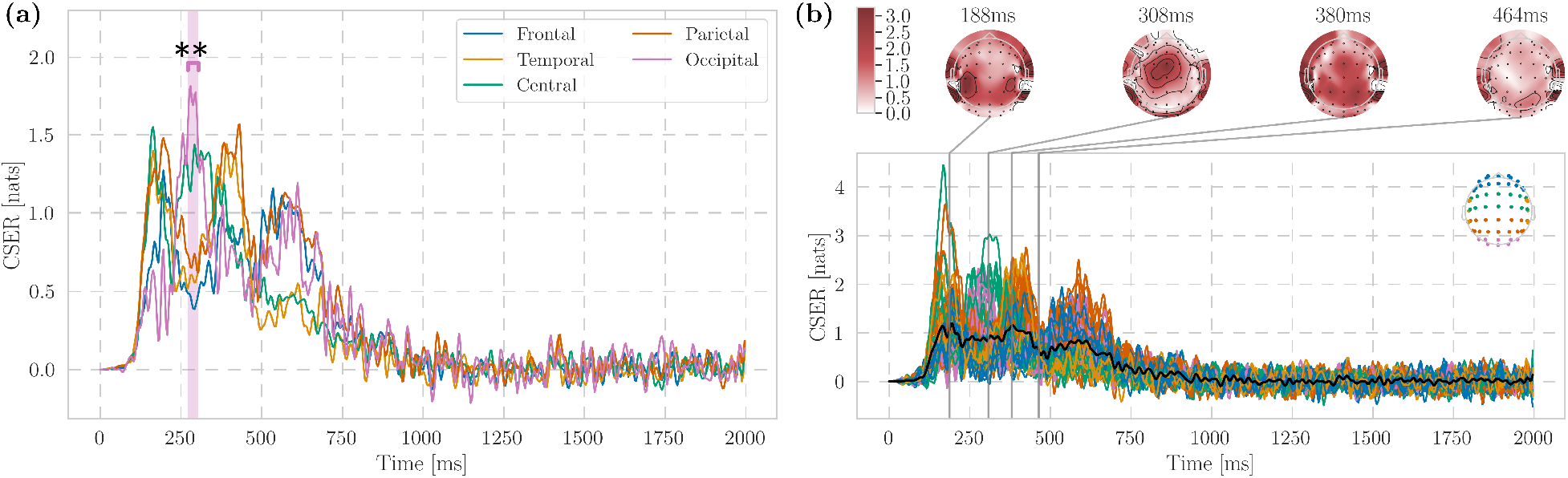
Time-resolved CSER shows a significant increase in the occipital region at approximately 300 ms. **(a)** Time-resolved CSER for different brain regions and averaged across subjects for *−*5 dB. **(b)** Time-resolved CSER for each channel and averaged across subjects for *−*5 dB. Topographic maps representing the overall complexity distribution are shown for specific time points. The CSER values shown are the differences w.r.t. CSER of pure noise. For visualisation purposes, the CSER signals above were low-passed to 50 Hz.

For all the analyses outlined above, analogous results are obtained using the audibility ratings as classification criteria instead of the SNR levels (Supplementary Material), linking our results to the subjective experiences of the participants.

### C. Information transmission

We complement the neural complexity analyses with an investigation of how entropy is shared across brain regions. Specifically, building on the state-space model introduced above, we calculated the mutual information rate (MIR), a standard information-theoretic quantity that captures the entropy rate shared by two processes over time, i.e. the rate of information transmission [55–57].

We fitted the SS model and computed MIR across brain areas on sliding windows of half a second post-stimulus, tracking how MIR evolves over time and space for the various SNR levels (Fig. 4a). The first notable result is that spatially-averaged MIR tends to be lower for higher SNRs (Fig. 4b), with SNR *−*5 dB being able to discriminate across all SNR levels (*p <* 0.01 under ANOVA cluster permutation test and pairwise cluster permutation test, Supplementary Material). Interestingly, MIR is lowest in the window between 400 and 900 ms, which corresponds to the first peak of the ERP (Fig. 2c), then becoming statistically indiscernible from the pre-stimulus baseline after approximately 1 s. Finally, we examined the spatial connectivity of time-averaged MIR between pairs of regions across SNRs, observing that the shared entropy rate gradually decreases with increasing SNR levels. This phenomenon is particularly pronounced in the temporal and parietal regions (Fig. 4c).

**FIG. 4:**
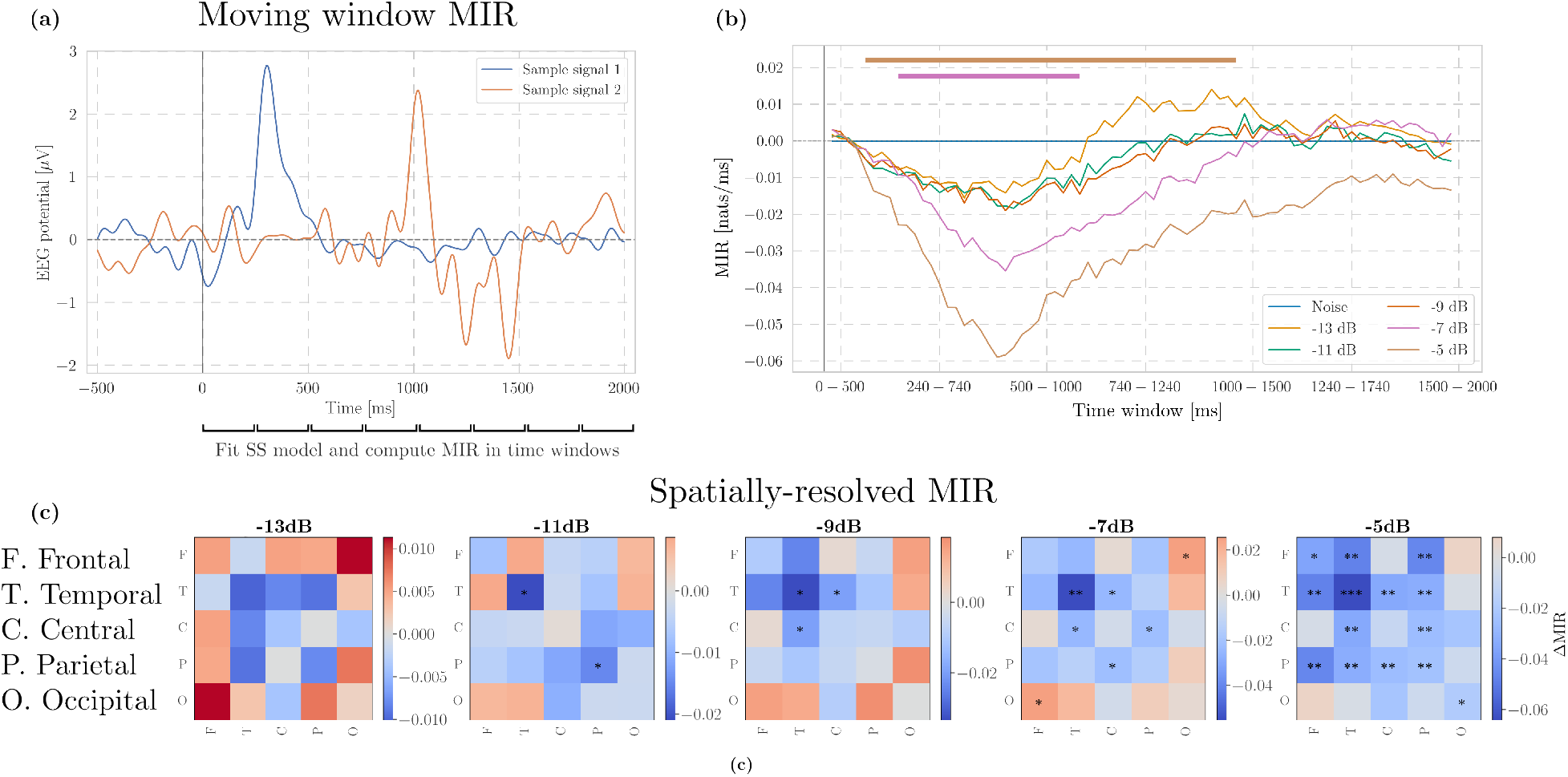
Mutual information rate decreases for higher SNRs. **(a)** We calculate mutual information rate (MIR) between two signals by fitting a separate SS model across sliding time windows. **(b)** Whole-brain averaged MIR decreases for clearer stimuli. Values shown are the differences w.r.t. MIR of pure noise. Coloured horizontal bars indicate significance level with *p <* 0.05 with a cluster permutation test. For visualisation purposes, only the statistically significant results against noise are shown. The full statistical analysis is reported in the Supplementary Material. **(c)** Spatial connectivity matrices of mutual information rate averaged across time for each SNR level. MIR shows a breakdown in information transmission, particularly localised in the temporal and parietal regions. The values shown are the differences w.r.t. MIR of pure noise. (P-values calculated with a one-sample t-test against the zero-mean null hypothesis. *: *p <* 0.05; **: *p <* 0.01; ***: *p <* 0.001).

## III. DISCUSSION

This study investigated how the organisation of information in the brain reflects progressive changes in perceptual awareness within a constant global state of consciousness. To this end, we examined neural complexity and inter-regional shared entropy rate in EEG activity using Complexity via State-space Entropy Rate (CSER) and the mutual information rate (MIR), in an auditory discrimination paradigm where stimulus clarity was systematically manipulated via signal-to-noise ratio. This approach allowed us to relate temporal and spectral complexity with perceptual clarity, and to examine corresponding changes in large-scale patterns of information transmission.

### A. Temporal complexity captures fine-grained changes in perceptual awareness

Time-resolved CSER revealed significant differences between non-perceptible stimuli and those ranging from the threshold of audibility to higher clarity, with the two clearest cues enabling discrimination across all perceptual levels and not only against noise. Specifically, higher levels of perceptual clarity were associated with increased neural complexity, with the most pronounced deviations from baseline broadly consistent with the timing of the N100 and P300 components in the ERP.

We propose that the observed increase in time-resolved complexity can be interpreted under two complementary perspectives. First, complexity may increase across the transition from unconscious to consciously accessible stimulus processing. In this view, greater complexity reflects the emergence of consciously differentiated content, indicating that when perceptual distinctions enter awareness, neural activity becomes more diverse and informative. Second, complexity may increase when perceptual content is experienced more clearly compared to when it is ambiguous. Here, the modulation of complexity reflects a graded aspect of awareness: as the clarity or confidence of perception increases, so might the local integration and differentiation of neural activity. Together, these hypotheses suggest that neural complexity may index both the presence and the richness of conscious perceptual content. These are two aspects that future work could systematically disentangle by designing paradigms that independently manipulate stimulus detectability and perceptual clarity.

### B. Neural complexity precedes and complements ERP responses

In our results, the first CSER peak precedes the ERP by approximately 30 ms. An analogous result was obtained in the original CSER study [21], in which time-resolved CSER was applied to ECoG data of a macaque undergoing an auditory oddball task. Taken together, these findings suggest that increases in neural complexity may reflect early stimulus-related differentiation in neural dynamics, potentially corresponding to the initial formation of prediction-error-like signals, which then shape the subsequent neural activity captured by the ERP.

Relatedly, CSER shows greater sensitivity than the ERP in distinguishing between perceptual states. This has twofold implications: firstly, it suggests that neural complexity captures aspects of stimulus processing that are not fully reflected in evoked responses alone. From a functional perspective, this may indicate that complexity measures provide complementary information about the temporal organisation of perceptual processing, underscoring the importance of the role of the prediction error in shaping neural dynamics – a link that future work could formalise by integrating temporal complexity measures with computational models of perception. Secondly, and independently of the former, these results have an operational application, as they provide a robust estimator that is more effective at distinguishing different perceptual experiences than the ERP, suggesting its potential as a sensitive marker of perceptual awareness in non-communicative or clinical populations where behavioural reports are unavailable.

### C. Markers of global states versus perceptual experiences

Much research has shown that neural complexity can characterise a variety of global states of consciousness, especially in terms of steady-state complexity and including well-known measures such as Lempel-Ziv complexity [9, 34]. In the present study, we found that spectrally resolved CSER revealed frequency-dependent changes in complexity. Higher SNRs and audibility were associated with lower CSER in the *α* and *β* bands, but higher CSER in the *δ* band, while CSER in the *θ* and *γ* bands did not differ significantly across conditions.

In contrast, we observed that broadband CSER failed to differentiate between SNR levels, and that this lack of discriminability persisted when the signal was localised to specific brain regions (Supplementary Material Fig. 6). Thus, given the extensive evidence that global complexity varies across states of consciousness, we suggest that this measure is better understood as a marker of the overall level of consciousness, rather than of the degree of perceptual experience. Accordingly, the lack of discriminability observed with steady-state broadband CSER likely reflects that participants maintained the same level of conscious awareness throughout the trials. This finding is also consistent with the indiscriminability across perceptual levels observed in the *γ* band, which aligns with previous work showing that complexity in the *γ* band is the main driver of broadband signal complexity [21, 42, 58, 59].

Time-resolved CSER, on the other hand, accurately discriminated between higher SNRs (−7 and *−*5 dB) and noisier stimuli. This highlights an important distinction between steady-state and time-resolved CSER: while steady-state complexity assumes stationarity of the signal, time-resolved CSER captures how complexity evolves over time, accounting for non-stationary neural dynamics. The superior sensitivity of time-resolved CSER in distinguishing perceptual levels underscores the importance of incorporating temporal fluctuations in the study of neural activity during cognitive tasks. How best to model these dynamics remains an open question, with ongoing debate regarding optimal approaches for capturing non-stationarities [21, 60–62].

### D. Global post-stimulus neural reorganisation in information transmission

The mutual information rate analysis revealed a global decline in information transmission for clearer auditory stimuli. Although there is a clear connection between complexity and predictability when looking at a single signal, for which one increases when the other decreases (see Sec. IV B, Eq. (13)), the present finding is distinct, as it involves pairs of brain regions at a time. Therefore, the breakdown of MIR elicited by the ERP suggests a rapid reorganisation in the information architecture of the brain, which is then re-established around 1 s post-stimulus, corresponding to the timing of the ERP decay. This decrease in shared entropy rate is mainly localised in the temporal and parietal regions. While the change within the temporal cortex can be readily understood as a consequence of auditory signal processing, the corresponding change in the parietal cortex is less expected. Within the context of auditory tasks, the parietal cortex is known to support attentional control [63, 64] and auditory–motor transformations involved in goal-directed actions and active reporting [65–68]. However, how these functional roles relate to the patterns of information transmission observed in our study remains unclear and warrants further investigation.

In sum, this investigation shows that changes in inter-regional shared entropy reflect both sensory processing and large-scale neural dynamics, revealing how auditory clarity modulates the brain’s global information architecture.

### E. Limitations and future work

An interesting line of research not explored in this study is the relationship between information dynamics in the brain and correct versus incorrect stimulus reports. Due to the skewed distribution of responses (with essentially no incorrect answers at higher SNRs), such an analysis was not feasible here. More generally, the analyses presented in this study are based on a single dataset and would thus benefit from further validation and generalisation across different paradigms. In this context, future research could extend the present investigations to other cognitive tasks, sensory modalities, and stimulus sets, contributing to a more comprehensive characterisation of neural complexity and the contents of consciousness.

On the methodological side, future work could focus on improving the data efficiency of CSER estimation. Because current methods require a careful balance among window length, sample size, and bias in parameter estimation, complexity estimates can be sensitive to the amount of available data and how it is segmented. Developing approaches that reduce this dependence – such as training models on longer pre-stimulus data to estimate single-trial complexity, as opposed to estimating it from the average of the residuals – would enable CSER to be applied at finer temporal and trial-level scales with greater reliability. Finally, given the relevance of non-stationary transients in ERP analyses, a promising avenue for future research involves exploring different models to include such dynamics. These include SS models with time-varying parameters or external non-stationary inputs [69], hidden Markov models [70], kernel-based approaches [71], and others [72].

## F. Conclusion

This work demonstrates how neural complexity can serve as a marker to discriminate between different percepts of awareness within the same global level of consciousness during an auditory discrimination task. By applying Complexity via State-space Entropy Rate (CSER), we found that broadband changes in perceptual awareness are reflected in variations of neural complexity that are temporally, but not spatially, localised. Furthermore, time-resolved CSER proved to be able to discriminate between dynamical structures across SNRs otherwise indiscernible by the ERP. On the other hand, the spectral decomposition of CSER revealed that conventional complexity markers, such as variations in the *γ* band, did not exhibit significant changes. Instead, a richer pattern was found, with increased complexity in the *δ* band and decreased complexity in *α* and *β*. Complementing these findings, the analysis of neural information transmission revealed both temporal and spatially specific reorganisations, consistent with the processing of the auditory cue.

Overall, these results highlight the importance of considering non-stationary dynamics and time-resolved estimators to link neural information processing and perceptual levels – underscoring the methodological difference in the study of global states of consciousness. We hope that this study encourages the application of similar methodologies to portray a richer taxonomy of information processing in cognitive tasks. We believe this avenue can deepen our understanding of brain function in cognitive processes across various aspects of perceptual experiences, while also providing robust estimators that can be employed in clinical settings.

## IV. METHODS AND DATA

### A. State-space models

State-space models are a class of statistical models that represent observed data as the noisy projection of an underlying latent process evolving over time. Thanks to their generality and flexibility, these frameworks have been widely applied in a broad range of areas, from econometrics [73, 74] and control theory [75, 76] to neuroscience [77, 78] and signal processing [79], and more recently in combination with information-theoretic tools [21, 80–82].

Following standard references of linear time series modelling [74, 83], here we provide a synthetic introduction to SS models before introducing CSER in the following section.

Given a time-varying signal *y*_*t*_, an SS representation of *y*_*t*_ is a mathematical model that describes its dynamics via a set of stochastic equations. More specifically, the SS representation of *y*_*t*_ involves a state equation that models the behaviour of an unknown state variable *ξ*_*t*_ (Eq. (1)), and an observation equation that relates the state variable with the actual result of the observations (Eq. (2)). These governing equations can be written as

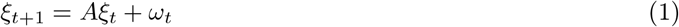

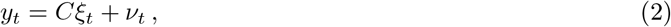

where *A, C* are matrices of coefficients, and *ω*_*t*_, *ν*_*t*_ are zero-mean white noise Gaussian terms sampled from

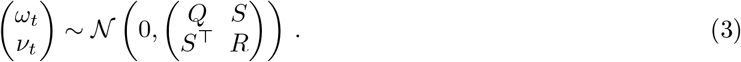

This framework accounts for the dynamics of *y*_*t*_ to emerge as a noisy observation of the hidden process *ξ*_*t*_ through the matrix *C*. On the other hand, the evolution of the state variable *ξ*_*t*_ is only determined by its previous state plus the noise term. This setup is particularly useful in real-data applications, in which the observed data is indeed a noisy measurement of an underlying process that cannot be directly gauged.

We can then introduce the quantity 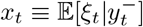 and the *innovation* term 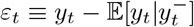, which can be shown to be i.i.d. white noise Gaussian terms with covariance *V*. Hence, using *x*_*t*_ and *ε*_*t*_, we can rewrite the SS equations in innovation form:

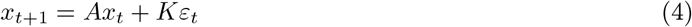

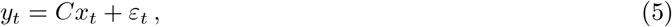

where *A, C* are the same matrices of Eqs. (1)-(2), and *K* is usually denoted as Kalman gain matrix. The advantage of innovation SS is that the noise terms are now the same in the state and observation equations, significantly simplifying mathematical derivations.

To determine the matrices *K, V* and fully obtain the innovation SS from the SS equations, we need to solve a Discrete Algebraic Riccati Equation (DARE) [84] that links the conditional variance of the state variable to the parameters of the model. In formula, this reads:

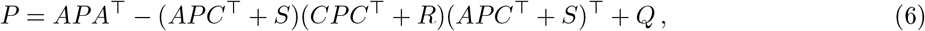

Where 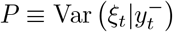. Once *P* is obtained, then *V* and *K* can be calculated as

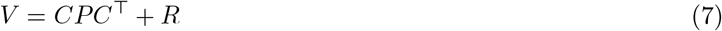

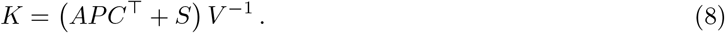

Hence, the knowledge of the parameters *A, C, K, V* completely determines the form of the (innovation) state-space. With a small abuse of notation, in the following we will indicate as DARE the procedure to obtain *V* from the SS parameters *A, C, Q, R, S*, so that we can write Eqs. (6)-(7) compactly as *V* = DARE (*A, C, Q, R, S*).

For practical purposes, solving the DARE of Eq. (6) can be avoided, as innovation SS models can be directly fit from data using e.g. state-subspace algorithms such as Larimore’s [79]. However, this treatment is necessary for outlining the computation of mutual information rate in the SS framework (Sec. IV C).

### B. Complexity via State-space Entropy Rate

Complexity via State-space Entropy Rate (CSER) is a measure designed to capture the complexity of dynamical systems by quantifying the predictability of their trajectories within a reconstructed state space [21]. By examining how the system evolves over time, CSER evaluates the rate at which statistical innovations are introduced in the system, reflecting the balance between order and randomness in the system’s behaviour. Overall, both steady-state and time-resolved CSER are particularly valuable for analysing non-linear and time-evolving systems, offering insights into their structural and stochastic properties and finding applications in fields such as neuroscience and complex systems [21]. Referring to the original work for a more exhaustive description of the method, here we provide a summary of how it operates, outlining how it can be implemented for this study.

A crucial ingredient for understanding CSER is the notion of state-space representation of a dynamical process introduced above. Starting from the formulation of the entropy rate of a stochastic process [85],

CSER builds upon the edifice of Gaussian SS models and estimates the dynamical complexity of the process *y*_*t*_ as the entropy of its innovation term:

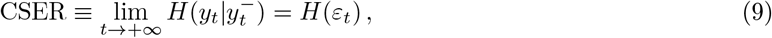

where the second equality directly follows from the definition of *ε*_*t*_ and the law of total expectation. Then, using the entropy formula for a Gaussian variable *X ~ 𝒩* (0, Σ)

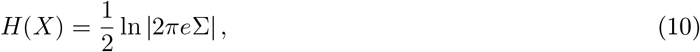

the expression of CSER simply reads

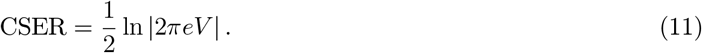

A convenient feature of state-space models is that they allow for a spectral decomposition of Eq. (11) as

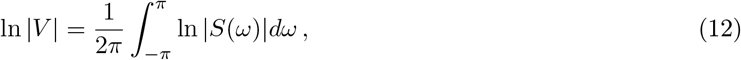

where *S*(*ω*) is the spectral density of the system, which can be directly evaluated via the transfer function of the system [83]. In other words, this formulation enables us to calculate how each frequency band of the signal *y*_*t*_ is contributing to the overall complexity of the process.

Moreover, an additional property of CSER is that it can be employed to compute instantaneous entropy rates, enabling it to track how the complexity of a stochastic process evolves over time. Given a time signal *y*_*t*_, this can be operationalised by fitting an innovation SS model on the baseline of *y*_*t*_, and then making one-step-ahead predictions using the parameters of the model. By evaluating the residuals between the predicted signal and *y*_*t*_, we can then estimate the residual covariance and compute time-resolved CSER as the residuals’ log-likelihood [21]. Hence, this procedure yields a local measure of entropy rate that describes how the real dynamics of *y*_*t*_ differ from what is expected from its baseline. This approach is particularly powerful for comparing trials representing different tasks, such as standard and deviant tones in an auditory oddball paradigm.

Finally, using Eq. (9), we remark here that CSER can be linked with time-delayed mutual information as

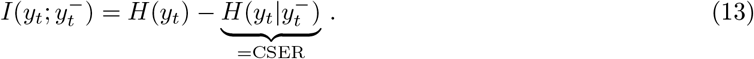

Since the variance of the process *y*_*t*_ is often normalised to 1 for numerical stability reasons, *H*(*y*_*t*_) can be treated as a constant, and an interesting property arises: up to some prefactors, complexity is the opposite of the information transmission over time. Taking into account this behaviour is essential to correctly interpret the results provided by CSER and MIR.

### C. Mutual information rate

Mutual information rate (MIR) is defined as the amount of information exchanged between two dynamical processes ***X, Y*** per unit of time. First introduced by Shannon [55] and then further formalised in [56, 57], the MIR between ***X*** and ***Y*** is defined as

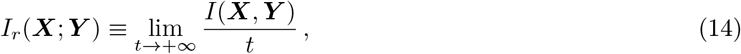

where ***X*** = (***X***_1_, …, ***X***_*t*_) and ***Y*** = (***Y***_1_, …, ***Y***_*t*_) are time series of length *t*, and *I*(***X, Y***) is the mutual information between ***X*** and ***Y***. This can be directly calculated via a combination of Shannon’s entropies as

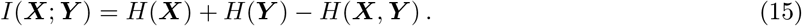

Putting together Eqs. (14)-(15) we obtain

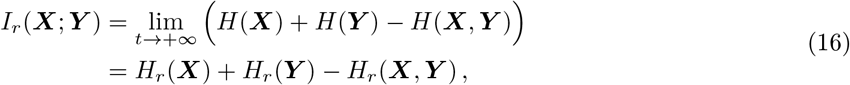

where we introduced the entropy rate *H*_*r*_(***X***) *≡* lim_*t→∞*_ *H*(***X***)*/t* [85].

Estimating MIR on large dynamical systems is in general non-trivial, as the size, complexity, and chaoticity of the system’s trajectories usually hinder the estimation of the probabilities needed for computing MIR [86]. However, calculating MIR on SS models avoids such issues as it takes advantage of the stationarity assumption underlying these frameworks. In fact, the entropy rate of a stationary process can be conveniently written as a conditional entropy in the limit of infinite future:

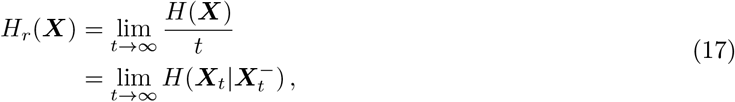

where we denoted with ***X***_*t*_, ***Y***_*t*_ the present states of the system, and with 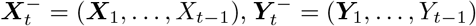 the full past of ***X***_*t*_ and ***Y***_*t*_, respectively. For a stationary process, MIR can then be computed from Eqs. (16)-(17) as

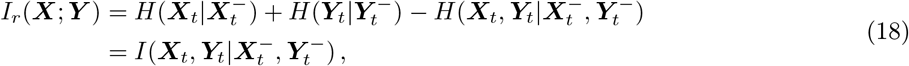

where we introduced the conditional mutual information *I*(*·*; *·*|*·*) [85].

Thus, given a SS model in innovation form (Eqs. (4)-(5))) and the corresponding parameters *A, C, K, V*, the entropy term *H* 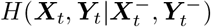 can be directly calculated using Eq. (10) and the conditional residual covariance *V*, which precisely specifies Var 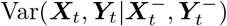. For the other two entropies of Eq. (18), the reduced systems of the full SS models are needed. To obtain this, we need to revert the innovation SS model into a canonical state space model, which is easily achieved by considering the term *η*_*t*_ ≡ *Kε*_*t*_ as a noise term on its own, so that in analogy with Eq. (3) we can write

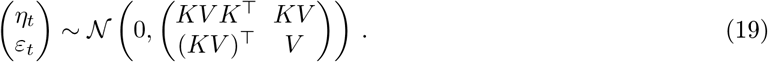

This allows us to solve the DARE (Eq. (6)) for a subset of the system and obtain the desired residual covariance conditioned on a part of the system. Specifically, if the dynamical variables have dimension *d*_*X*_ and *d*_*Y*_, i.e. 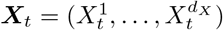 and 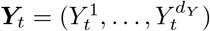, then the full innovation SS parameters in the observation equation can be written in blocks as

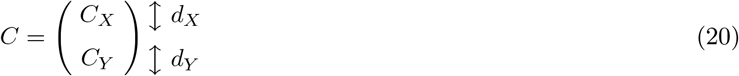

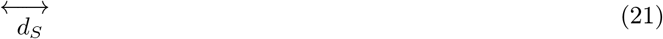

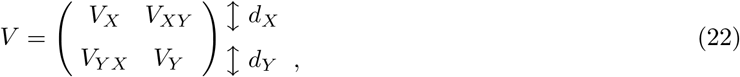

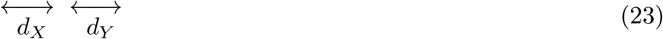

where *d*_*S*_ is the dimension of the state variable.

Following a similar derivation to that of Sec. IV A, this notation allows us to solve the following DAREs to find the residual covariance of the reduced system:

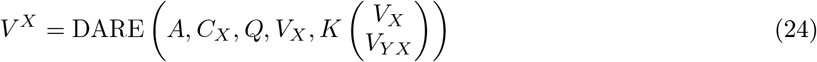

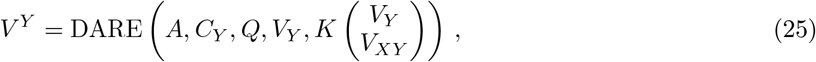

where we used the upper index to discriminate 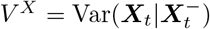 from 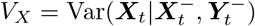.

Hence, we can calculate the remaining conditional entropies of Eq. (18) using *V* ^*X*^, *V* ^*Y*^, and Eq. (10), then finally compute MIR.

### D. Neural data

The data we considered for the analysis is the dataset collected by Sergent *et al*. [52], comprising EEG recordings from 20 adult participants (10 women and 10 men, mean age 23.4 years, range 21–29), all of whom were right-handed native French speakers. The experiment involved auditory stimuli presented in the form of French vowels (“a” and “e”) embedded in threshold equalising noise. These vowels were presented at five different signal-to-noise ratios (SNRs): −13, −11, −9, −7, and −5 dB, alongside “noise-only” trials. Each stimulus lasted for 200 ms and was followed by random inter-trial intervals. Participants completed 160 trials for each condition, for a total of 960 trials per session.

The experiment was conducted across two sessions (active and passive), performed on different days and counterbalanced in order. During the active session, participants identified the vowels and rated their subjective audibility on a scale from 0 to 10. In the passive session, participants listened to the same stimuli without task requirements, engaging instead in unrelated tasks designed to probe mind-wandering or response speed. In this work, we specifically focused on the active session, as it ensured clearer distinctions in perceptual clarity across stimuli.

EEG data were recorded using a Brain Products system with 64 active electrodes arranged in the 10-20 system, referenced to FT10. The continuous EEG signal was sampled at 500 Hz, with a high-pass filter set at 0.003 Hz during acquisition. Preprocessing involved high-pass filtering at 0.4 Hz, manual artefact rejection, independent component analysis (ICA) for removing eye movement artefacts, and re-referencing to the average signal across electrodes. Data were epoched around stimulus onset (−500 to 2000 ms) and baseline-corrected using the pre-stimulus interval. Approximately 94% of trials per participant were retained after artefact removal. In this work, we performed our analysis directly on the preprocessed data.

For a more comprehensive description of the dataset, we refer to the original study [52].

### E. Neural analysis

We applied the methods outlined above to analyse the neural signal of the 6 different SNR levels (Sec.II) and the audibility ratings (Supplementary Material) in the auditory discrimination task considered, focusing on the active session. For the spatially localised analyses, we investigated the frontal, temporal, central, parietal, and occipital regions, which contain the following electrodes:

**TABLE 1:**
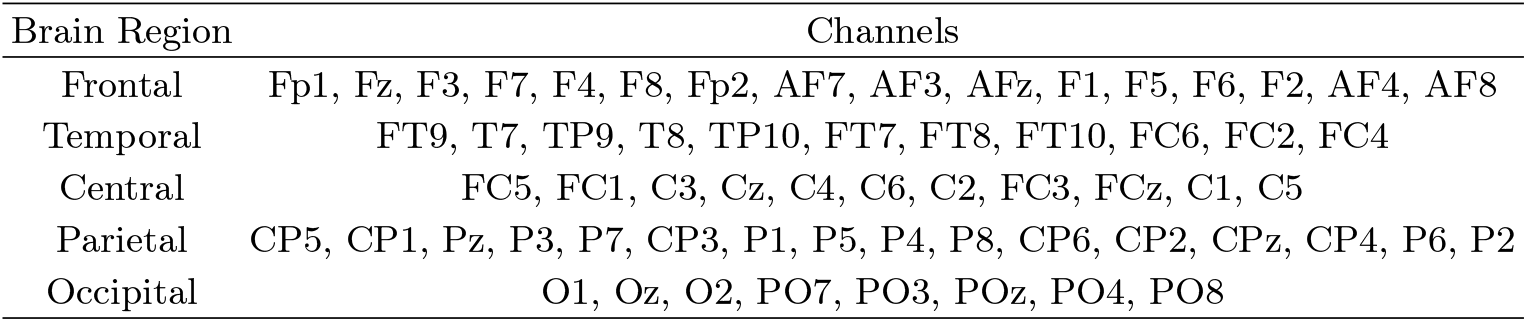
EEG channels grouped by brain region.

| Brain Region | Channels |
| --- | --- |
| Frontal | Fp1, Fz, F3, F7, F4, F8, Fp2, AF7, AF3, AFz, F1, F5, F6, F2, AF4, AF8 |
| Temporal | FT9, T7, TP9, T8, TP10, FT7, FT8, FT10, FC6, FC2, FC4 |
| Central | FC5, FC1, C3, Cz, C4, C6, C2, FC3, FCz, C1, C5 |
| Parietal | CP5, CP1, Pz, P3, P7, CP3, P1, P5, P4, P8, CP6, CP2, CPz, CP4, P6, P2 |
| Occipital | O1, Oz, O2, PO7, PO3, POz, PO4, PO8 |

We first investigated the overall complexity of the signals via the steady-state CSER approach. First, we divided the EEG data into six groups, each corresponding to different SNR levels and containing multiple trials. For each electrode, we fit a state-space model to the set of post-baseline time series of each group, obtaining the SS parameters *A, C, K, V* for each of them. Using these parameters, we compute CSER as per Eq. (11). We then repeated this process for all electrodes and subjects, resulting in 64 CSER values per subject, each corresponding to a complexity estimate for a specific electrode. We followed an analogous approach for the calculation of the spectrally-decomposed quantities, in which we computed CSER via Eq. (12) for each frequency band (*δ ∈* (1, 4) Hz, *θ ∈* (4, 8) Hz, *α ∈* (8, 12) Hz, *β ∈* (12, 30) Hz, *γ ∈* (30, 100) Hz).

As for the time-resolved CSER, we followed a slightly different strategy. As above, we began by selecting an electrode and grouping its trials into 6 categories, each with a corresponding SNR. For each group, we then fit a state-space model only on the baseline of the time series, and used the parameters of the model to make one-step-ahead predictions. Hence, we computed the residuals as differences between the predictions and the real time series, which we then averaged before taking the log-likelihood. Repeating this procedure for each electrode and participant, we obtain a set of time series that describe how neural complexity evolves in time for each brain area of all subjects. Analogous procedures apply for the computation of CSER on the different auditory ratings.

For the calculation of mutual information rate, we split each participant’s data into the 6 SNR groups as explained above, and then fit the SS model and computed MIR on sliding windows of length 0.5 s. This allowed us to retain enough data points for an accurate estimation of the SS model, while keeping the assumption of signal stationarity reasonable.

## Acknowledgments

We would like to deeply thank Tom Froese, whose support and precious insights were indispensable to the realisation of this project.

## Appendix A: Additional results

### 1. Single-trial ERP and CSER

The fact that time-resolved CSER faithfully tracks the neural activation in the ERP is not straightforward. In particular, the SS model from which CSER is computed is not trained on the average baselines of Fig. 2c, but on multi-trial data in which patterns are hard to identify. This can be seen explicitly in Fig. 5, where all the trials for SNR *−*5 dB of subject 2 are plotted, together with their mean (black). Unlike in Fig. 2c, the behaviour of the baselines does not seem entirely detached from the dynamics of the ERP, as the size of the baseline fluctuations with those of the ERP is comparable. This fundamental property is then entirely lost when an average is performed, leading to an oversimplification of the dynamics of the system. Hence, although seemingly intuitive, it is highly non-trivial that the complexity of the signal should correlate with its amplitude.

**FIG. 5:**
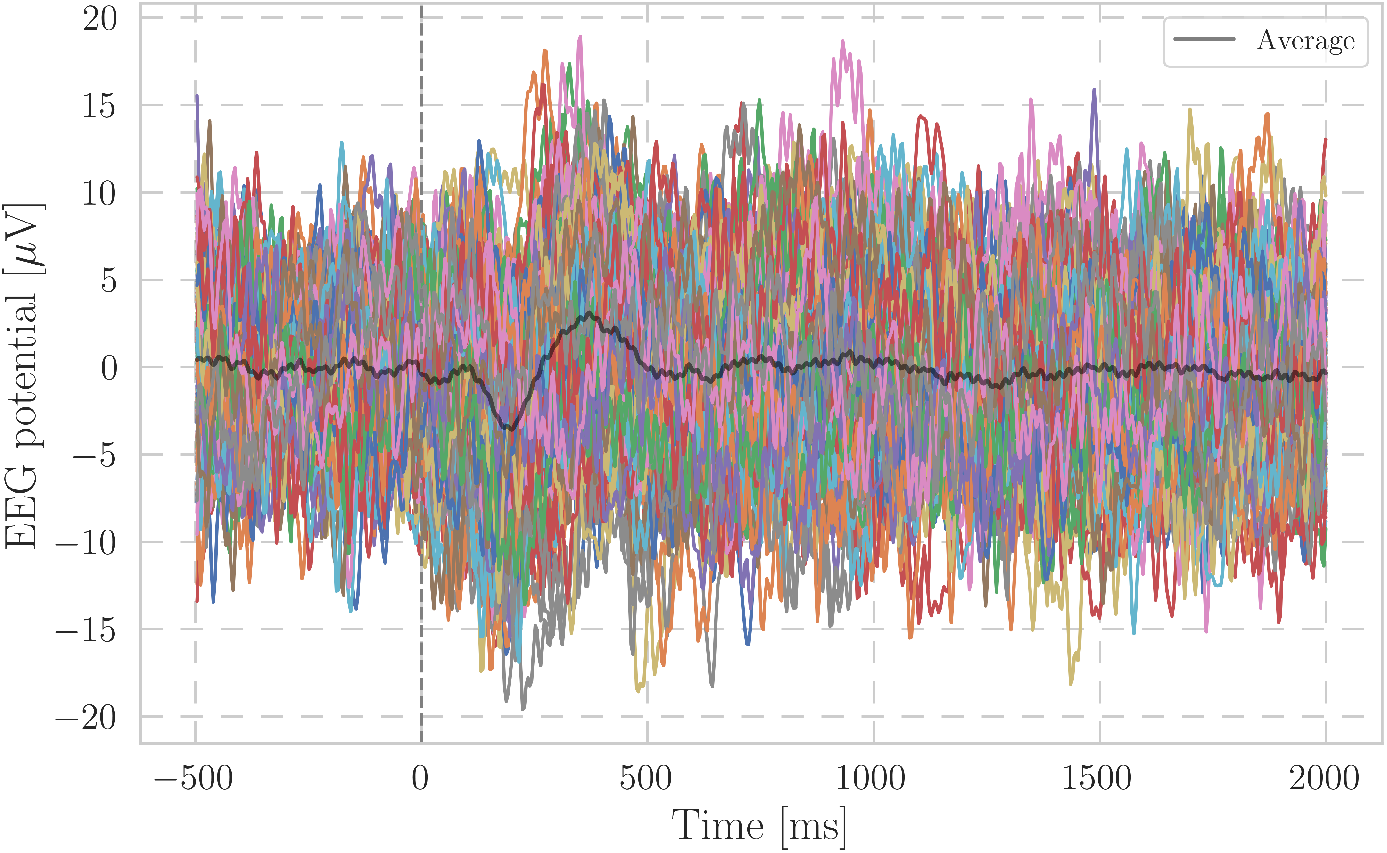
ERP across all trials for SNR *−*5 dB and subject number 2. The black line represents the average.

**FIG. 6:**
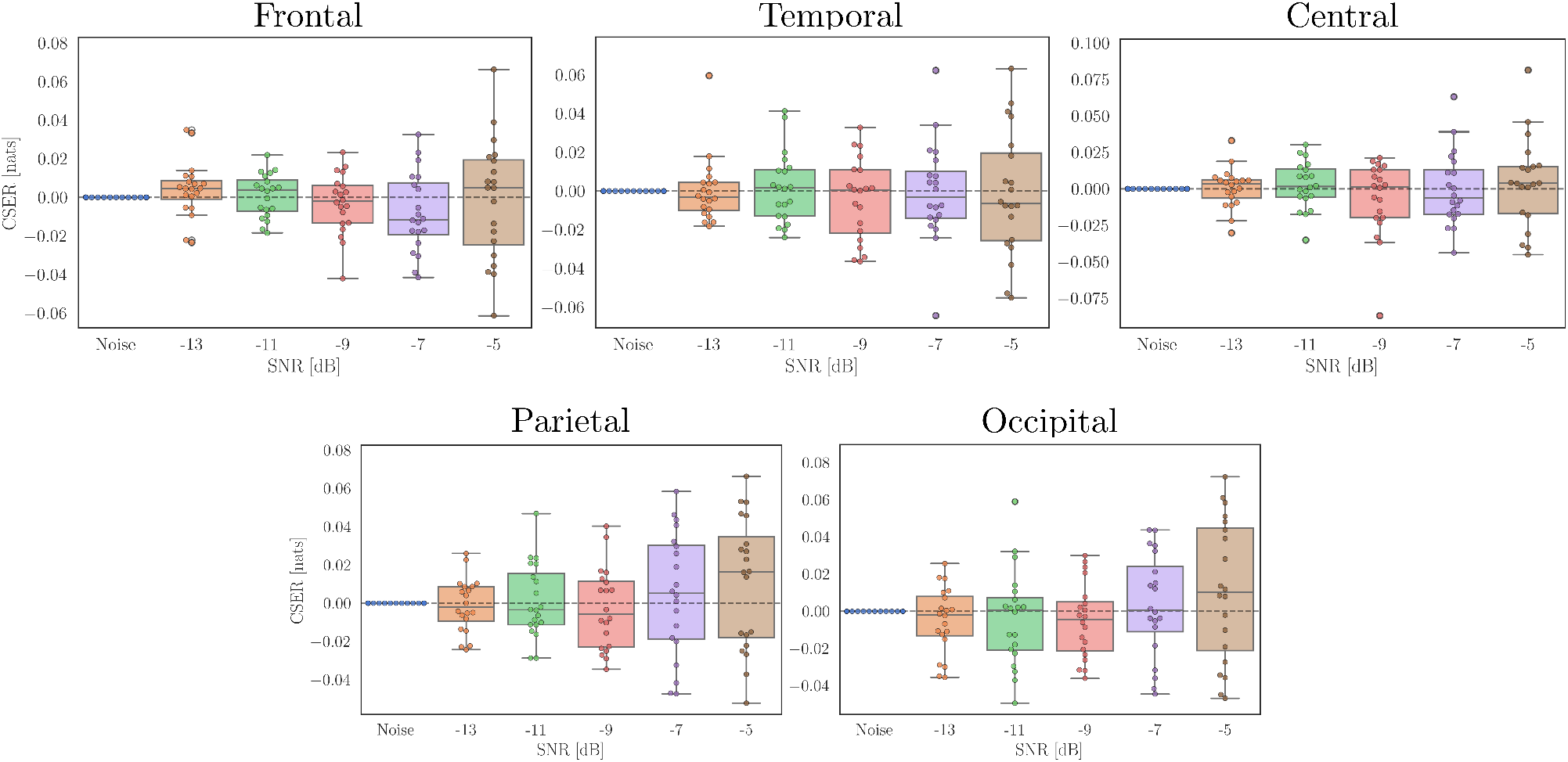
Steady-state CSER for frontal, temporal, central, parietal, and occipital brain regions for different SNR levels and all subjects. The CSER values shown are the differences w.r.t. CSER of pure noise.

**FIG. 7:**
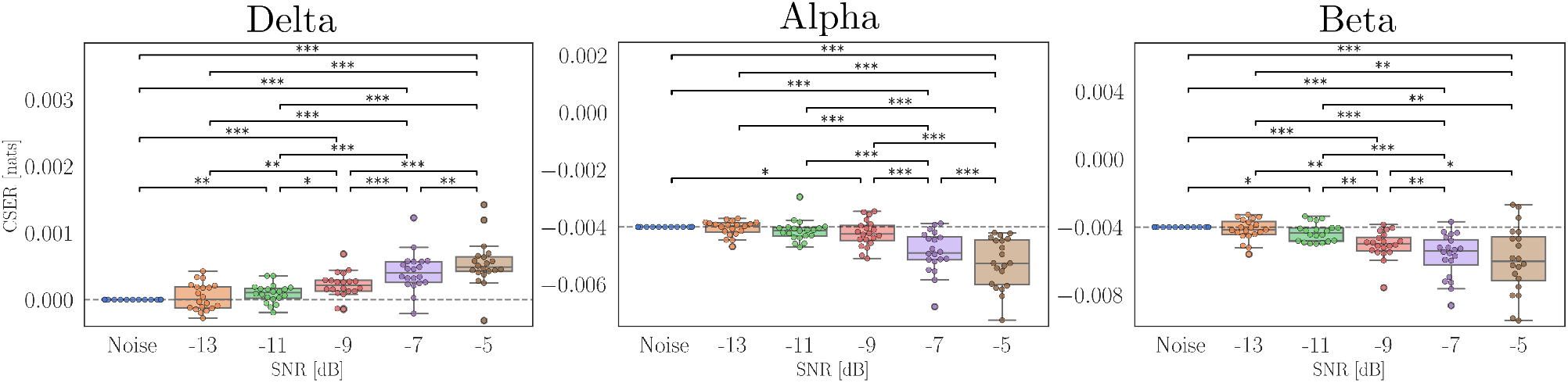
Complete statistical results for *δ, α*, and *β* frequencies of steady-state CSER across SNR levels. Values represent differences relative to CSER obtained from pure noise and are averaged across all subjects. Statistical significance was assessed using one-sample t-tests against zero (null hypothesis of no deviation from noise), followed by Holm–Bonferroni correction. Asterisks denote corrected significance levels (*: *p <* 0.05; **: *p <* 0.01; ***: *p <* 0.001). Only significant post-hoc comparisons are shown.

**FIG. 8:**
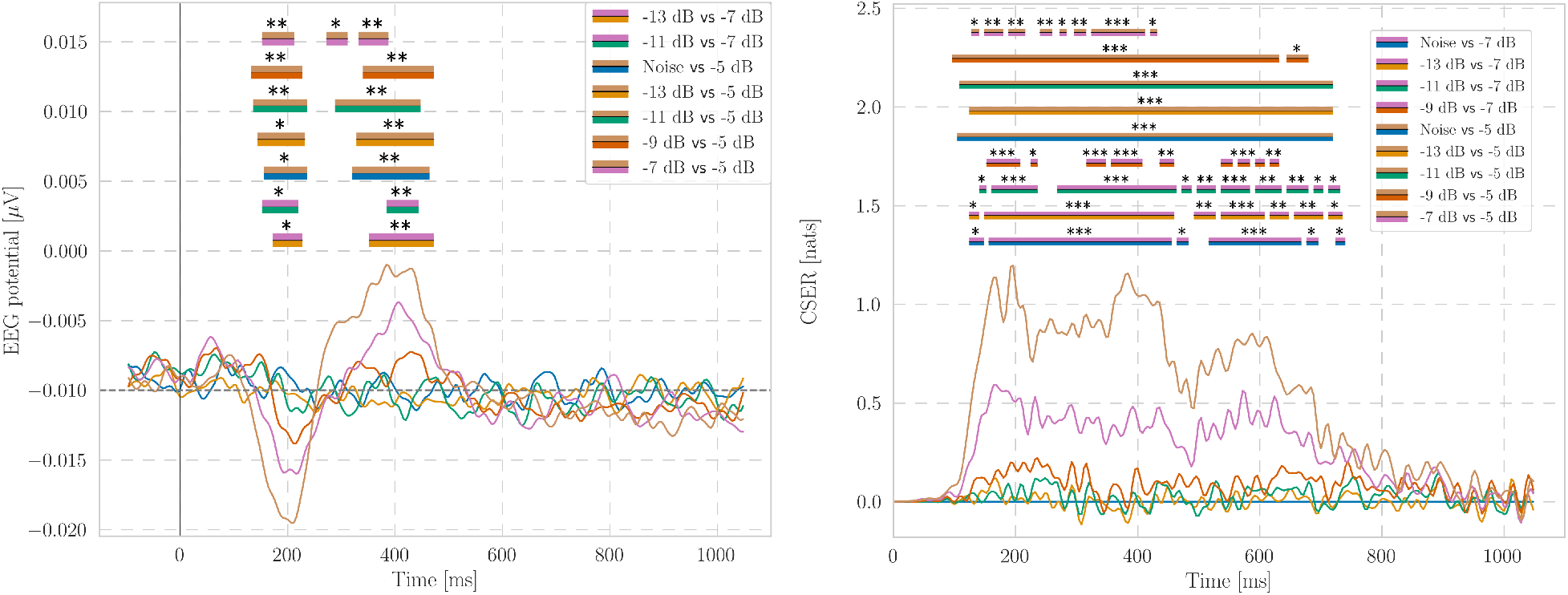
Complete statistical results for time-resolved analyses. Pairwise post-hoc comparisons between SNR levels are shown for ERP and time-resolved CSER, averaged across all channels and subjects. CSER values are expressed as differences with respect to pure noise. Statistical significance was assessed using cluster-based permutation testing, and all p-values were Holm–Bonferroni corrected (*: *p <* 0.05; **: *p <* 0.01; ***: *p <* 0.001). Only comparisons surviving correction are displayed.

**FIG. 9:**
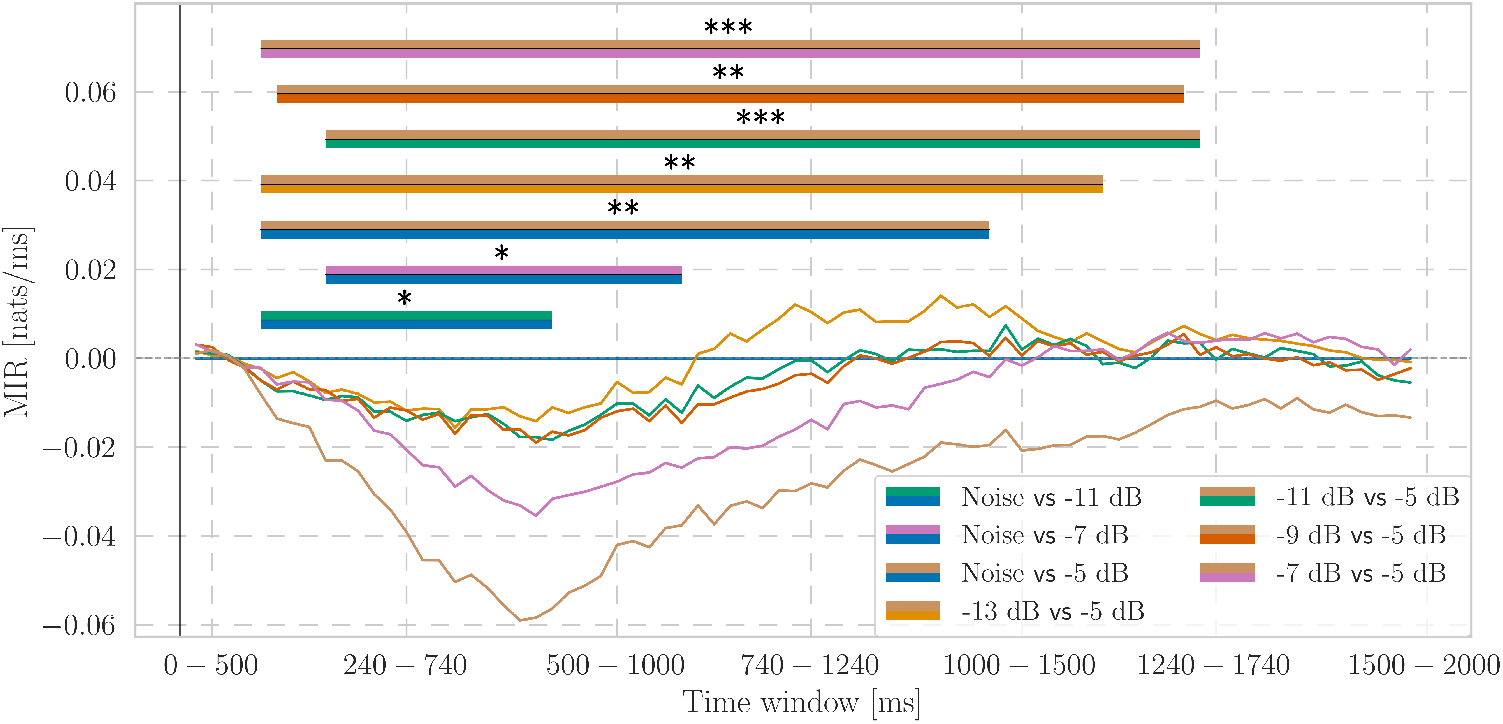
Complete statistical results for time-resolved whole-brain averaged MIR across SNR levels. MIR values are expressed as differences relative to pure noise. Statistical significance was determined using cluster-based permutation tests with Holm–Bonferroni correction for multiple comparisons (*: *p <* 0.05; **: *p <* 0.01; ***: *p <* 0.001). Only pairwise comparisons surviving correction are displayed.

### 2. Spatial steady-state CSER

We complement the analysis of steady-state CSER of the main text by showing that even a more fine-grained analysis with a subdivision in different brain regions does not show any significance between the complexity of various SNRs (Fig. 6).

### 3. Further statistical analyses

In this section, we report the complete statistical results corresponding to the main analyses presented in the paper. Specifically, we provide all statistically significant pairwise post-hoc comparisons. All reported p-values were corrected for multiple comparisons using the Holm-Bonferroni method.

For time-resolved analyses (ERP, steady-state and time-resolved CSER, and MIR), statistical significance was assessed using non-parametric cluster-based permutation tests [54]. For steady-state CSER analyses, significance was evaluated using one-sample t-tests against a zero-mean null hypothesis. Only effects surviving correction are displayed.

### 4. Audibility ratings

In addition to the investigations based on different SNR levels, here we present analogous studies considering audibility ratings as markers of experience. Since during the experiments participants were asked to rank the audibility rating on a scale from 0 to 10, here we group the answers into three categories: Low (0, 1, 2, 3, 4), Medium (5, 6, 7), and High (8, 9, 10) audibility. Applying the procedures above on this threefold classification, similarly to the SNR scenario, we obtain that time-resolved CSER accurately tracks the potential activations in the ERP (Fig. 10).

**FIG. 10:**
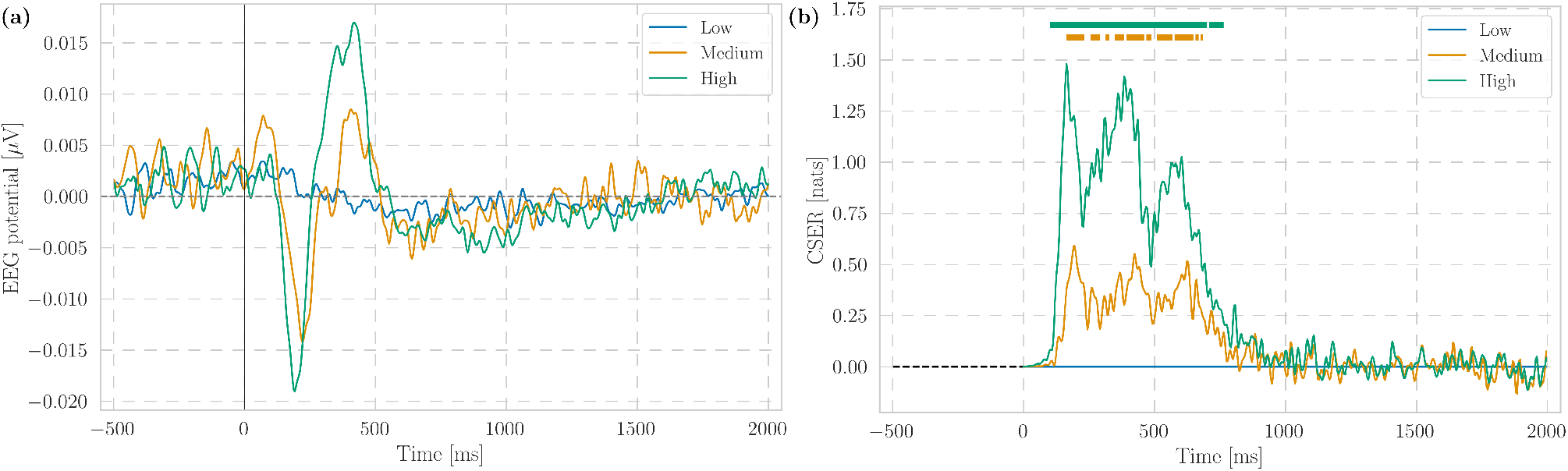
(a) EEG ERP divided by audibility ratings levels averaged across all channels and subjects. (b) Time-resolved CSER divided by audibility ratings and averaged across all channels and subjects. The CSER values shown are the differences w.r.t. CSER of Low audibility. For visualisation purposes, the CSER and EEG signals above were low-passed to 50 Hz.

On the other hand, computing steady-state CSER and its decomposition on the full time series, we observe the same patterns: statistically higher complexity in *δ* and lower in *α, β*, but no overall broadband difference between the ratings, confirming the results above (Fig. 11).

**FIG. 11:**
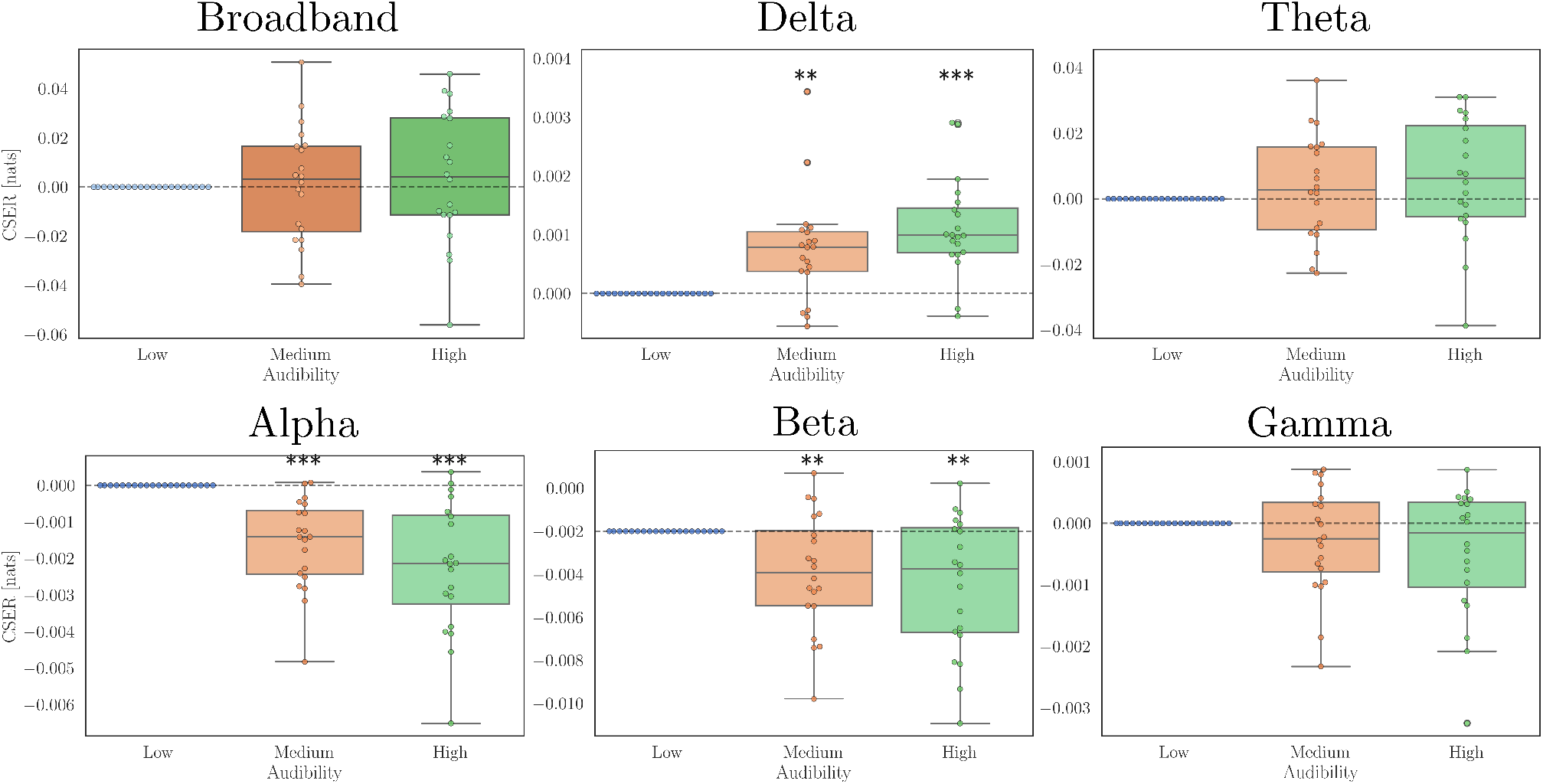
Broadband and frequency-decomposed steady-state CSER for different audibility ratings and for all subjects. The CSER values shown are the differences w.r.t. CSER of Low audibility. (P-values calculated with a one-sample t-test against the zero-mean null hypothesis. *: *p <* 0.05; **: *p <* 0.01; ***: *p <* 0.001).

These findings should not come as a surprise, as there is a strong correlation between High audibility ratings and SNR −5 dB trials, and Low audibility and SNRs of pure noise, −11, −13 dB.

